# Changes in the blood plasma lipidome associated with response to atypical antipsychotic treatments in schizophrenia

**DOI:** 10.1101/2020.03.25.008474

**Authors:** Valéria de Almeida, Guilherme L. Alexandrino, Adriano Aquino, Alexandre F. Gomes, Michael Murgu, Paul C. Guest, Johann Steiner, Daniel Martins-de-Souza

## Abstract

Atypical antipsychotics are widely used to manage schizophrenia symptoms. However, these drugs can induce deleterious side effects, such as MetS, which are associated with an increased cardiovascular risk to patients. Lipids play a central role in this context, and changes in lipid metabolism have been implicated in schizophrenia’s pathobiology. Furthermore, recent evidence suggests that lipidome changes may be related to antipsychotic treatment response. The aim of this study was to evaluate the lipidome changes in blood plasma samples of schizophrenia patients before and after 6 weeks of treatment with either risperidone, olanzapine, or quetiapine. Liquid chromatography tandem mass spectrometry (LC-MS/MS) analysis showed changes in the levels of ceramides (Cer), glycerophosphatidic acids (PA), glycerophosphocholines (PC), phosphatidylethanolamines (PE), phosphatidylinositols (PI), glycerophosphoglycerols (PG), and phosphatidylserines (PS) for all treatments. However, the treatment with risperidone also affected diacylglycerides (DG), ceramide 1-phosphates (CerP), triglycerides (TG), sphingomyelins (SM), and ceramide phosphoinositols (PI-Cer). Moreover, specific lipid profiles were observed that could be used to distinguish poor and good responders to the different antipsychotics. As such, further work in this area may lead to lipid-based biomarkers that could be used to improve the clinical management of schizophrenia patients.

## 1. Introduction

Schizophrenia is a chronic mental disorder characterized by positive, negative, and cognitive symptoms (Owen et al., 2016). Currently, antipsychotics are the most common form of treatment for schizophrenia. Typical antipsychotics were the first-generation developed to treat the symptoms of schizophrenia, but can lead to extrapyramidal side effects. The negative impact that these side effects had on health drove the search for other options, which are known as atypical antipsychotics. Clinical data reported superiority of atypical compared to typical antipsychotics in treatment of the negative and cognitive symptoms (Owen et al., 2016). However, atypical antipsychotics also presented some side effects, such as metabolic syndrome (MetS) and obesity, which are associated with increased cardiovascular disease risk for schizophrenia patients. Thus, the metabolic effects of atypical drugs could be associated with the premature mortality observed in schizophrenia (Laursen et al., 2012; Mitchell et al., 2013). However, it is not clear whether patients with schizophrenia are predisposed to metabolic disorders or if these disorders are a consequence of the treatment (Fernandez-Egea et al., 2009).

Atypical antipsychotics present different profiles on metabolic disturbances. For instance, the dibenzodiazepines clozapine and olanzapine have a greater effect on metabolic processes compared to other atypical drugs, such as the dibenzothiazepine quetiapine or other classes that include risperidone and aripiprazole (Raben et al., 2017; Spertus et al., 2018). Although the biological mechanisms have not been completely elucidated, some studies have suggested that metabolic disturbances promoted by atypical drugs could be due to effects on histamine type 1 (H1), serotonin 2 C (5HT2C), and muscarinic acetylcholine type 3 (M3) receptor systems in central and peripheral systems (Hahn et al., 2011). Therefore, differential pharmacological targets of the antipsychotics could explain the diversity in metabolic effects, including on lipid metabolism.

Lipids are widely found in the brain and play a role in membrane composition, energy metabolism, neurotransmission, and neuromodulation of multiple molecular pathways. Lipids in cellular membranes can modulate the functions of certain integral proteins (Piomelli et al., 2007). As modulators, some lipids, such as phosphatidylinositol, can be involved in signaling cascades that affect several neuronal and glial functions (Piomelli et al., 2007). Therefore, alterations in metabolic and structural lipids may potentially be associated with the pathophysiology of disorders such as schizophrenia (Misiak et al., 2017). Particularly, phospholipids have a crucial role in the structure of neuronal membranes and appear to be associated with the pathobiology of schizophrenia (Horrobin, 1998). Furthermore, studies have suggested that changes in lipids can be associated with an increased breakdown of membrane phospholipids, and pro-inflammatory state, suggesting that the modulation of lipid homeostasis is a potential target for the improvement of schizophrenia symptoms (Keshavan et al. 2003; Leppik et al. 2019; Wood, 2014, 2019). Thus, lipidomic changes may have a potential for disease-state or predictive biomarkers for schizophrenia, or a predictive lipidic signature for antipsychotic response in schizophrenia (Aquino et al., 2018).

Considering that disturbances in lipid pathways have been found in schizophrenia, this study aimed to investigate lipid alterations promoted by atypical antipsychotics and their association with good or poor response. The blood plasma lipidomic profiles of drug-naïve schizophrenia patients were investigated before and after treatment with either quetiapine, olanzapine or risperidone using shotgun, liquid chromatography, tandem mass spectrometry (LC-MS/MS).

## 2. Methods

### 2.1 Patients and antipsychotic treatment

We recruited 54 patients – including male and female – who fulfilled the Diagnostic and Statistical Manual (DSM)-IV criteria for schizophrenia. The ages of patients ranged from 16 to 66 years. Patients were treated for 6 weeks with either risperidone (n=23), olanzapine (n=17) or quetiapine (n=14) (Table S1). Patients were grouped as good or poor responders to the medication, based on whether or not a 50% reduction of total Positive and Negative Syndrome Scale (PANSS) and Structured Clinical Interview (SCID-I) (First et al., 2002) scores occurred. Exclusion criteria included substance abuse, symptoms induced by a non-psychiatric medical illness or treatments. Blood samples were collected between 8 and 9 AM (fasted) by venous puncture in the psychiatric clinic at the University of Magdeburg, Germany. All procedures were approved by the Institutional Review Board of the University of Magdeburg (process 110/07, from November 26^th^, 2007 amended on February 11th, 2013). All patients signed an informed consent document prior to their participation in the study.

### 2.2 Plasma sampling and Sample Preparation

Blood samples were collected by venous puncture and plasma was prepared from patients at baseline (T0) and after the 6-week treatment period (T6) at the psychiatric clinic at the University of Magdeburg. Lipids were extracted from 25 μL plasma by addition of 100 μL isopropanol, vortexing 1 min, and then incubating 10 min at room temperature. The samples were kept at −20°C for 18 h and centrifuged at 7.500g for 20 min. The supernatant was collected, dried in a vacuum centrifuge, and reconstituted in 100 μL of buffer (50% isopropyl alcohol, 25% acetonitrile, and 25% water) prior to LC-MS/MS analysis (Sarafian et al., 2014).

### 2.3 LC-MS/MS

All chemicals were purchased from Sigma-Aldrich (Seelze, Germany) and were high performance liquid chromatography (HPLC) purity grade. An Acquity H-Class UPLC system (Waters Corporation, Milford, MA, USA) was used as the inlet for mass spectrometry. The column was an ACQUITY UPLC^®^ CSH C18, 2.1 x 100 mm, 1.7 μm particle size (Waters Corporation), operating at a flow rate of 0.4 mL/min at 55°C. The chromatographic separation was performed in gradient mode using a mobile phase system consisted by two solvents, A and B. Solvent A was ACN:water (60:40, v:v) containing 10 mM ammonium formate and 0.1 % formic acid, and phase B was IPA:ACN (90:10, v:v) containing 10 mM ammonium formate and 0.1 % formic acid. Gradient steps started with 60% A and 40% B, changing linearly to reach 57% A and 43% B at 2.0 min, 50% A and 50% B at 2.1 min, 46% A and 54% B at 12.0 min, 30% A and 70% B at 12.1 min, attaining a final composition of 1% A and 99% B at 18.0 min, before returning to the initial composition of 60% A and 40% B to compose a total chromatographic stage of 18.1 minutes. The buffer remained at this composition until 20 min prior to preparing the system for the next injection. The injection volume was 1.0 μL.

The inlet system was coupled to a hybrid Xevo G-2 XS quadrupole orthogonal time-of-flight mass spectrometer (Waters Corporation, Manchester, UK), controlled by MassLynx 4.1 software. Data were acquired in positive electrospray ionization (+)-(ESI) mode with the capillary voltage set at 2.0 kV, the cone voltage at 30 V, and the source temperature at 150°C. The desolvation gas was nitrogen (N^2^), with a flow of 900 L/h at 550°C. Data were acquired from 100 to 2000 mass/charge (m/z) in MS^E^ mode, during which the collision energy was alternated between low (2 V) and high (ramped from 20 to 30 V). Leucine-enkephalin [C_28_H_37_N_5_O_7_, ([M+H]^+^ = *m/z* 556.2771)] (Waters Corporation) was used as lock mass reference at 0.2 ng/L at flow rate of 10 μL/min.

### 2.4 Data processing and software

Progenesis QI 2.2 software (Nonlinear Dynamics, Indianapolis, IN, USA) was used to process data for peak selection. For compound identification, the Lipidmaps database was used with a precursor mass error ≤ 10 ppm and fragment tolerance ≤ 10 ppm. Other parameters such as fragmentation score were also considered for disambiguation. The whole dataset contained 1584, 1604, and 1584 normalized m/z values per patient sample for the risperidone, olanzapine, and quetiapine treatments, respectively. Multivariate data analysis was performed to search in the pooled m/z values for the signals responsible for any significant differences between pre- and post-drug treatment, using m-file functions in the Matlab R2016a (Matworks, Natick – MA, USA) and the Pls Toolbox 8.11 (Eigenvector Research Inc., Wenatchee, WA, USA).

### 2.5 Multivariate data analysis

The dataset belonging to patients from each treatment were imported into Matlab as single data matrices **X*i***_(IxJ)_, where *i* = 1 (risperidone), 2 (olanzapine), or 3 (quetiapine); I = number of samples in the corresponding treatment (this corresponds to each pre- and post-drug treatment samples per patient); J = number of m/z signals common to all samples of the same treatment. The submatrices **G*i*** and **B*i*** built from **X*i*** containing only responders or non-responders, respectively for the *i-th* treatment, were analyzed similarly to **X*i***, searching for m/z signals specific to treatment effect.

### 2.6 ML-PLSDA

The main differences in the overall profile of the m/z values between pre- and post-drug treatment in the patients within each treatment were assessed by ML-PLSDA. Multilevel statistical approaches consider the paired structure of the subjects in a temporal or longitudinal study, in which all subjects act as their own control. This allows greater ability to distinguish any within- and between-subject variances in the original dataset. This is usually done in univariate statistics using paired t-test and repeated measures ANOVA, but a similar approach can also be extended to multivariate data (van Velzen et al., 2008). In ML-PLSDA, the within-level variance data were initially extracted from **X*i*** to the new matrix **X_W_*i***_(I, J)_, according to previous study (Westerhuis et al., 2010). A variable selection approach based on variable importance in projection (VIP) scores was performed in **X_W_*i*** within ML-PLSDA, using double cross-validation (2CV) (Szymańska et al., 2012), to obtain only those m/z values significant to the treatment effect (i.e., distinguishing the pre- (Class 1) and post- (Class 2) drug treatment patient samples). The 2CV-ML-PLSDA procedure is described as follows: **X_W_*i*** was randomly split into 2 submatrices **X_W_*i***_CAL(A, J)_ and **X_W_*i***_TEST(B, J)_, A = 32, 24 or 16, and B = 12, 8 or 6, for the risperidone, olanzapine and quetiapine treatments, respectively. ML-PLSDA was performed only in the **X_W_*i***_CAL_, removing approximately 25% of the samples during the venetian blind (inner) cross-validation for model optimization. During this cross-validation process, **X_W_*i***_CAL_ was arranged to extract row pairs from the same patients (i.e., pre- and post-drug treatment), obtaining the correct number of latent variables (LVs) for the models in the lowest root mean squares error of cross-validation (RMSECV). After model optimization, only the VIP scores upon a threshold were selected while extracting the most informative m/z signals for the classification in **X_W_*i***_CAL_. This threshold was defined iteratively to the point below which the ML-PLSDA models did not increase their RMSECV. This was computed for the new model every time a new threshold was established. The overall 2CV procedure was performed up to 500 iterations for non-repetitive combinations of the samples in **X_W_*i***_CAL_ and **X_W_*i***_TEST_, and only those m/z signals that were selected in at least 85% (risperidone) or 60% (olanzapine and quetiapine) of the iterations were considered relevant to distinguish the treatment effects in the patients. The remaining m/z signals in each **X_W_*i***_CAL_ were removed as their presence resulted in an overall increase of the RMSECV and misclassifications while predicting **X_W_*i***_TEST_ after another round of iterations performed similarly. This combined variable selection and 2CV-ML-PLSDA approach provided the most important (and generalizing) m/z signals from the pooled data without fitting overly-optimistic models, given that the final m/z signals were selected based exclusively on non-biased histograms from random iterations. Additionally, the model predictions were correctly evaluated, because they were computed for totally independent samples [i.e., when the samples belonged only to the test set (**X_W_*i***_TEST_)]. A similar approach to this work for biomarker identification using PLSDA has been published elsewhere (Khakimov et al., 2016). The 2CV-ML-PLSDA approach was also performed separately for the **G*i*** (responders) and **B*i*** (non-responders) matrices accordingly.

### 2.7 One-way ANOVA F-test and Principal Component Analysis

The final m/z signals selected after ML-PLSDA were individually ranked according to the magnitude of their changes during each treatment, using one-way ANOVA F-test (α ≤ 0.01). Furthermore, the principal component analysis (PCA) comprising the auto scaled, selected m/z signals from the patients in each treatment group highlighted the multivariate patterns of the corresponding biomarkers most associated with each treatment.

## 3. Results

### 3.1 Clinical features

According to the criteria set in the Methods section, 18, 12, and 5 patients showed a good response to risperidone, olanzapine, and quetiapine treatment, respectively (Table 1). There was no significant effect for BMI before (T0) and after 6-weeks (T6) for quetiapine (*t*(13) = −1.31, p = 0.2139), olanzapine (*t*(16) = −2.11, p = 0.0512), while a significant effect was observed in risperidone-treatment (*t*(22) = −3.54, p = 0.0018).

**Table 1:** Characteristics and clinical measurements of patients

|  | **Risperidone** | **Olanzapine** | **Quetiapine** |
| --- | --- | --- | --- |
| **Patients (n)** | 23 | 17 | 14 |
| **Good response (%)** | 78 | 70 | 36 |
| **First episode (%)** | 35 | 41 | 43 |
| **Age (mean ± SD)** | 39 ± 13 | 37 ± 11.2 | 34 ± 8.6 |
| **Female/Male (n)** | 6/17 | 9/8 | 7/7 |
| **BMI T0 (mean ± SD)** | 24.7 ± 4.6 | 26.9 ± 4.1 | 23.8 ± 3.7 |
| **BMI T6 (mean ± SD)** | 25.9 ± 4.2 | 27.7 ± 3.7 | 24.2 ± 4.6 |
| **Cigarette smoking (%)** | 65 | 70 | 79 |

BMI - body mass index; Good response - 50% reduction of total PANSS

### 3.2 Selection of m/z signals associated with response

The Multilevel Partial Least Squares – Discriminant Analysis (ML-PLSDA) models showing the m/z signals most associated with the treatments (Table 2). The low CVs and prediction misclassifications throughout the random 2CV iterations infer non-biased robustness and relevance of the selected m/z signals as potential classifiers of the effects of the treatments in the patients, given that the models correctly classified most of the patients before and after treatment. The one-way ANOVA F-tests performed for each selected m/z rejected the null hypothesis, i.e. the m/z signals were not significantly different across the treatments. The visual separation between patients before and after treatment with risperidone, olanzapine, or quetiapine is depicted in Figure 1 to clarify that the m/z signals obtained by 2CV-ML-PLSDA were affected by the antipsychotic treatments. After PCA on the data containing only the biomarkers selected by the 2CV-ML-PLSDA approach, the first principal component therefore expresses the main source of variance distinguishing pre-versus post-treatment patients that was selected from the multi-level modeling (Figure 2). However, the investigations of the next PCs did not reveal any clear separation between good and poor responders that could also come from the selected biomarkers. Therefore, even regarding the m/z signals corresponding to lipids most affected by the treatments, there is no clear evidence in the data that the lipids affected by the treatments were also affected differently between good and poor responders. This is an indication that there may be other compounds affecting specifically good or poor responders, requiring local, multi-level modeling using only good or poor responders to potentially extract these target compounds.

**Table 2.** Number of selected m/z signals from the pooled dataset, CVs, and prediction misclassifications after obtaining the most important m/z signals (metabolites) related to pre- and post-drug treatments for schizophrenia, using 2CV-ML-PLSDA analysis.

|  | **Selected**<br>**m/z** | **CV-misclassification\*<sup>1</sup>**<br>**(%)** | **Pred. misclassification\*<sup>2</sup>**<br>**(%)** |
| --- | --- | --- | --- |
| **Risperidone** |  |  |  |
| All Patients | 32 | 0.16 (1.86) | 0.35 (2.72) |
| Good Responders | 22 | 0.74 (3.50) | 0.92 (5.02) |
| Poor Responders | 47 | 1.7 (4.1) | 0 |
| **Olanzapine** |  |  |  |
| All Patients | 13 | < 0.01 (0.19) | 0.65 (2.78) |
| Good Responders | 13 | 1.30 (5.56) | 0.20 (1.82) |
| Poor Responders | 7 | 0 | 0 |
| **Quetiapine** |  |  |  |
| All Patients | 17 | 0.65 (3.90) | 0.30 (2.22) |
| Good Responders | 9 | 0 | 0 |
| Poor Responders | 5 | 5.0 (11.2) | 0 |

\* Standard deviation in the parentheses.
<sup>1</sup>CV errors while optimizing ML-PLSDA models in 2CV interactions.
<sup>2</sup> Class-label errors while predicting test sets in 2CV interactions.

**Figure 1.**
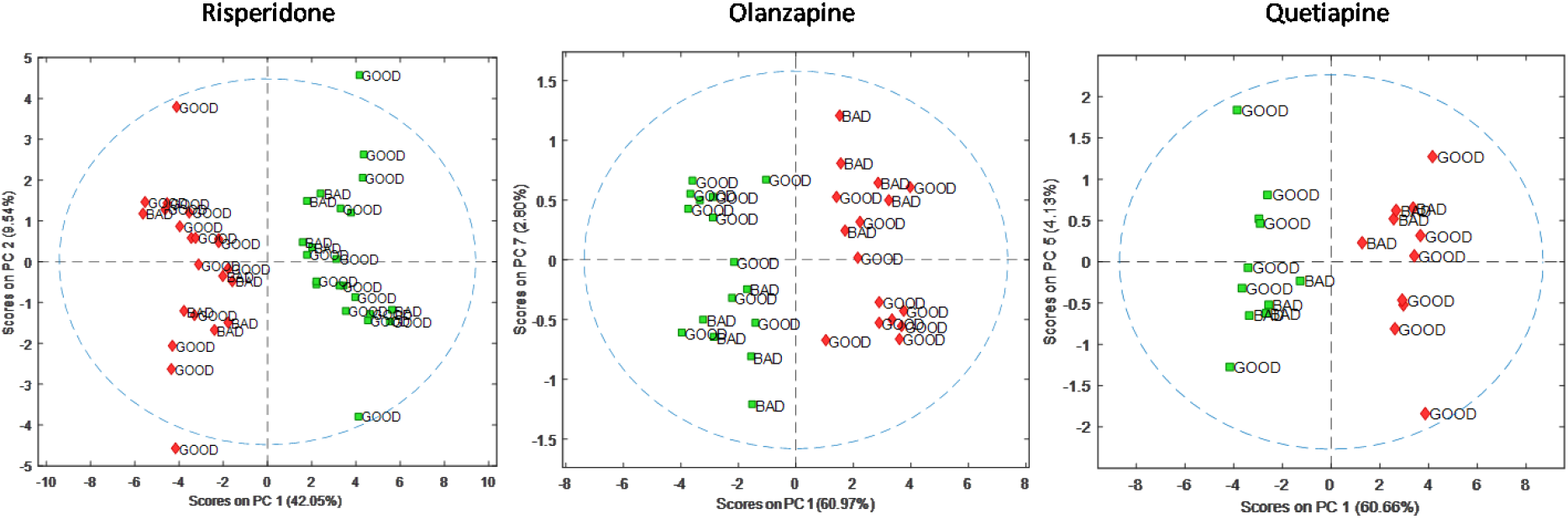
PCA scores plot distinguishing patients before (Green) and after (Red) treatments according to their respective profile of the m/z signals extracted by 2CV-ML-PLSDA. The samples were numbered continuously in the plot. The PC numbers in the y-axis express the best correlation and distinguish good and poor responders using the selected m/z signals during each treatment.

**Figure 2.**
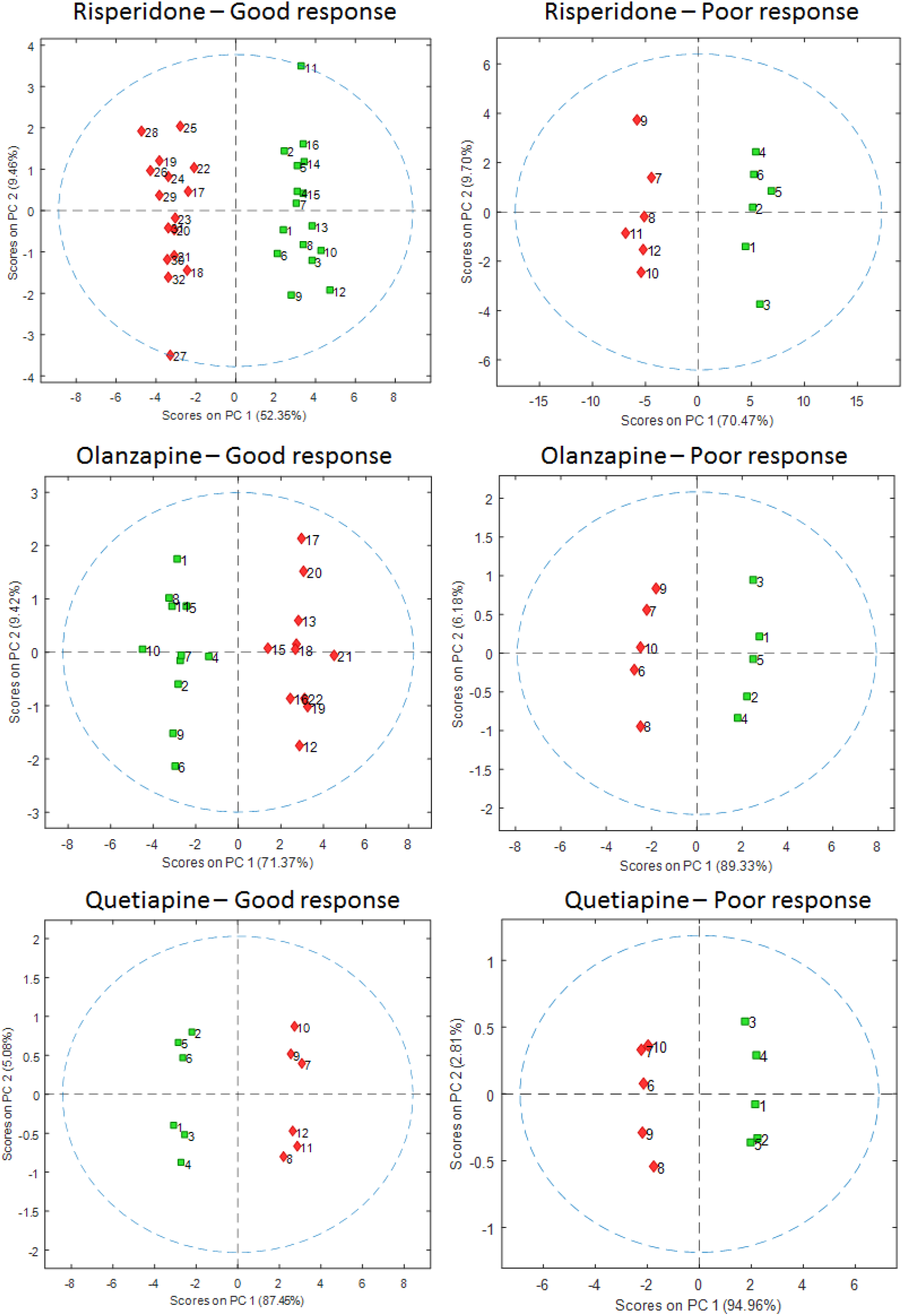
PCA score plot demonstrating the selected m/z signals from 2CV-ML-PLSDA distinguishing good and poor responders before and after the individual treatments.

### 3.3 Lipidomic changes for good and poor response groups

The main effects on lipids associated with their respective changes in poor or good responders of risperidone, olanzapine, and quetiapine are demonstrated in Tables 3, 4, and 5, respectively. Risperidone affected several lipid classes in the good and poor responder groups. Olanzapine affected mainly the PS, PC, PA, and PG lipid classes. Lastly, quetiapine affected the lipid profile of patients to a smaller extent.

**Table 3:** Risperidone effects associated with good or poor response

|  | PG (m/z 831.5516) | PC (m/z 508.3747) | PS (m/z 840.5693) | PS (m/z 812.5225) | SM (m/z 837.6211) | CerP (m/z 668.4981) | PA (m/z 645.4850) | PS (m/z 824.5750) | DG (m/z 613.4787) | PS (m/z 794.5280) | PC (m/z 868.5270) | PC (m/z 870.5429) | TG (m/z 907.7158) | PC (m/z 874.5758) | PC (m/z 808.5834) | PC (m/z 880.7094) | PI-Cer (m/z 892.5660) | PC (m/z 800.6515) | PC (m/z 832.5791) | PE (m/z 692.5198) |
| --- | --- | --- | --- | --- | --- | --- | --- | --- | --- | --- | --- | --- | --- | --- | --- | --- | --- | --- | --- | --- |
| Poor responders | ↓ | ↓ | ↓ | ↓ | ↓ | ↓ | ↓ | ↓ | ↓ | ↓ | ↓ | ↓ | ↑ |  |  |  |  |  |  |  |
| Good responders |  |  |  |  |  |  |  |  |  |  |  |  |  | ↓ | ↓ | ↓ | ↓ | ↑ | ↑ | ↑ |

**Legend:** CerP, Ceramide 1-phosphates; DG, diacylglycerides; PC, glycerophosphoglycerols; PE, phosphatidylethanolamines; PG, glycerophosphoglycerols; PI, phosphatidylinositols; PS, phosphatidylserines; SM, sphingomyelins; LarCer, lactosylceramides; TG, triacylglycerides. $p < 0.05$ , pre- compared to post-treatment

**Table 4:** Olanzapine effects associated with good or poor response

|  | PS (m/z 813.5488n) | PS (m/z 771.5408n) | PG (m/z 1578.1972) | PA (m/z 825.5735) | PG (m/z 1578.1972) | PC (m/z 832.5791) | PA (m/z 703.5610) |
| --- | --- | --- | --- | --- | --- | --- | --- |
| Poor responders | ↓ | ↓ | ↑ | ↓ |  |  |  |
| Good responders |  |  |  |  | ↑ | ↑ | ↑ |

**Legend:** PA, glycerophosphatidic acids; PC, glycerophosphoglycerols; PG, glycerophosphoglycerols; PS, phosphatidylserines. $p < 0.05$ , pre- compared to post-treatment

**Table 5:**
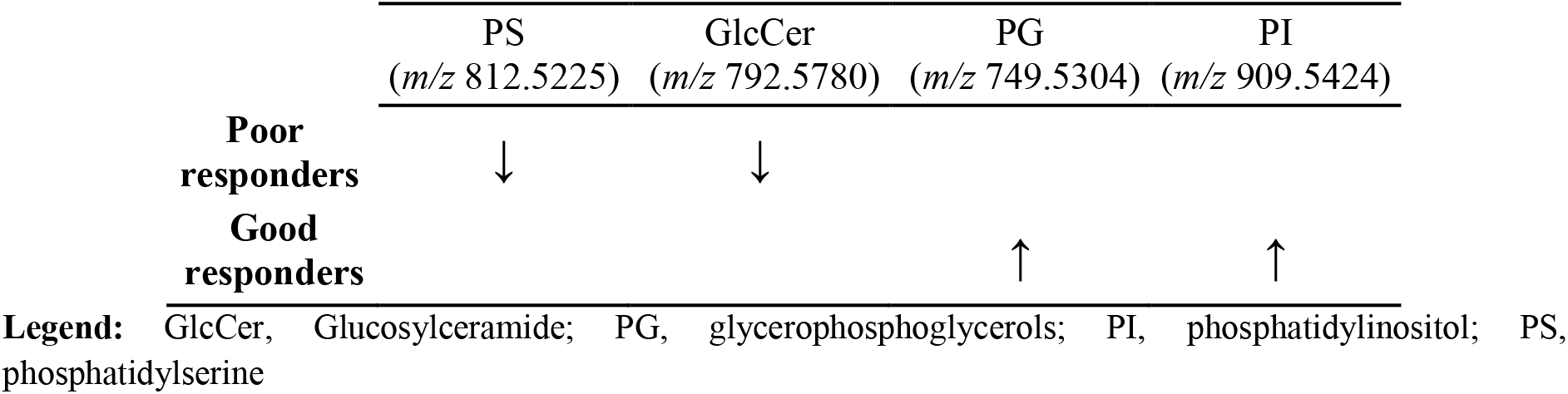
Lipid changes in poor or good responder patients treated with quetiapine

|  | PS<br>(m/z 812.5225) | GlcCer<br>(m/z 792.5780) | PG<br>(m/z 749.5304) | PI<br>(m/z 909.5424) |
| --- | --- | --- | --- | --- |
| Poor responders | ↓ | ↓ |  |  |
| Good responders |  |  | ↑ | ↑ |

**Legend:** GlcCer, Glucosylceramide; PG, glycerophosphoglycerols; PI, phosphatidylinositol; PS, phosphatidylserine

## 4. Discussion

Atypical antipsychotics have been widely used to treat schizophrenia; however, these drugs have been associated with the MetS, including dyslipidemia and obesity. Atypical antipsychotic drugs are diverse in chemical structure, and consequently the varying molecular mechanisms lead to differences in therapeutic response and side effects. Studies have reported that clozapine and olanzapine are associated with pronounced changes in lipids, whereas quetiapine and risperidone have moderate effects on lipids, while mild or no changes occur following aripiprazole-treatment (Koro et al., 2002; Lauressergues et al., 2011; Nasrallah et al., 2016). Accordingly, we found that olanzapine affected a higher number of lipid classes compared to quetiapine treatment. Conversely, risperidone affected more lipid classes compared to the other antipsychotics, in apparent contrast to a head-to-head meta-analysis that reported olanzapine had more impact on metabolic parameters compared to quetiapine and risperidone (Rummel-Kluge et al., 2010). A possible explanation for the above-mentioned discrepancies is that long-term treatment used in the meta-analyses, while the effects evaluated in this study were the result of a shorter period.

According to the literature, there are at least two different pathways in MetS promoted by atypical antipsychotics: first, studies have shown that atypical drugs modulate the functions of the hypothalamic nucleus, leading to increase in appetite (Kaur and Kulkarni, 2002); second, peripheral mechanism changes in the metabolism of glucose causing conditions such as insulin resistance, among others (Raben et al., 2017). Regarding lipid metabolism, atypical antipsychotics increase the sterol regulatory element-binding protein type 1 (SREBP1). This protein which regulates the expression of genes involved in the synthesis of cholesterol, fatty acid, triacylglycerol, and phospholipids (Bertolio et al. 2019), that we found affected by all antipsychotics investigated here (Table with classification). SREBP1 is found in several tissues, but the increase of this protein in adipose tissue and the liver seems to be the main factor of the dyslipidemia promoted by antipsychotics (Hellard et al., 2009; Lauressergues et al., 2011). SREBP1 also regulates lipid biosynthesis in the brain, and it can play a key role in the pathophysiology of schizophrenia or in the therapeutic effects of atypical drugs (Chen et al., 2017). Additionally, SREBF1 and SREBF2 genes, which encoded SREBP1 and SREBP2, were associated with genetic risk factor for schizophrenia and as target of antipsychotics as reviewed by Steen and colleagues (Steen et al. 2017).

The diversity in dyslipidemia promoted by atypical antipsychotics may be explained by the different mechanisms of action of these drugs. Olanzapine has been associated with several mechanisms of action, while fewer mechanisms have been reported for risperidone and quetiapine. As cholinergic (M3), histamine (H1), and serotoninergic (5HT2C) receptors are potential targets for MetS promoted by atypical antipsychotics, it is of interest that olanzapine and quetiapine are thought to act on these receptors (Sato et al., 2015), but not risperidone or its metabolite paliperidone. Thus, the changes in lipids promoted by olanzapine and quetiapine observed in this study may be related to these properties. However, the changes occurring after the treatment with risperidone cannot be explained in the same manner. However, it is expected that some antipsychotic mechanisms in MetS, along with behavioral patterns, such as poor diet, sedentary lifestyle, and smoking, are key contributors for MetS and cardiovascular risk (Sowden and Huffman, 2009). Additionally, ethnicity, age, and gender can modify the cardiovascular risk in patients with atypical treatment (Mitchell et al., 2013).

Studies have reported increased levels of triglycerides (TG) by atypical antipsychotics (Kaddurah-Daouk et al., 2007; Lally et al., 2013; Procyshyn et al., 2007; Sharma et al., 2014). Elevation of TG levels is probably the earliest and most sensitive index of the metabolic abnormalities associated with antipsychotics (de Leon et al., 2007). However, we observed that only risperidone treatment increased TG levels. In agreement with our findings, others studies have also reported hypertriglyceridemia in schizophrenia patients treated with risperidone (Kaddurah-Daouk et al. 2007; Khalili et al., 2007; Weinbrenner et al., 2009). Yet another study reported that only olanzapine out of the remaining atypical drugs promoted elevation of TG levels (Atmaca et al., 2003). Interestingly, atypical drug-induced hypertriglyceridemia has been associated with an improvement in symptoms of schizophrenia (Atmaca et al., 2003; Lally et al., 2013; Procyshyn et al., 2007; Sharma et al., 2014). Conversely, we found that an elevation in TG occurs in poor responders, suggesting that this effect is not associated with better outcomes in the schizophrenia symptoms as observed in those studies. Therefore, we suggest that hypertriglyceridemia seems not to be a good biomarker for a good response to antipsychotic treatment, and it could be associated with MetS, consequently increased cardiovascular risk. Additionally, we found a significant increased body mass index (BMI) – another component of MetS – of the patients after 6-week treatment with risperidone (before and after 6-week treatment), and a significance trend for increased BMI on olanzapine group. Accordingly, studies have shown weight gain with risperidone and olanzapine (Lauressergues et al., 2011; Musil et al., 2015; Raben et al., 2017).

Phosphatidylserine (PS) class was uniquely associated with a poor response in all treatments. We found decreased levels of PS (38:1, 36:2, O-38:2, and S-38:1) for risperidone, PS (38:3, and 36:3) for olanzapine, and PS (O-36:2, P-36:2) for quetiapine. PS lipids are widely found in neurotransmitter vesicle and axonal button membranes, and their main function described in the brain is related to cognition and attention processes (Zhang et al., 2015). Specifically, PS lipids are involved in dopamine release in the brain (Matam et al., 2016). Thus, we hypothesized that the antipsychotic-induced decrease in PS (Figure 3) could be associated with cognitive deficits in schizophrenia, resulting in worse outcomes in some components of the PANSS scale due changes in the dopamine release. This hypothesis is strengthened by the fact that PS reversed the reserpine-induced memory impairments in an animal study (Alves et al. 2000). Although the results show that PS lipids might be a potential biomarker for poor response to antipsychotics, further investigations are needed to confirm this hypothesis. In addition, PS lipids also can act as a messenger in apoptosis processes (Matam et al., 2016), suggesting a possible role in dopaminergic synapses.

**Figure 3.**
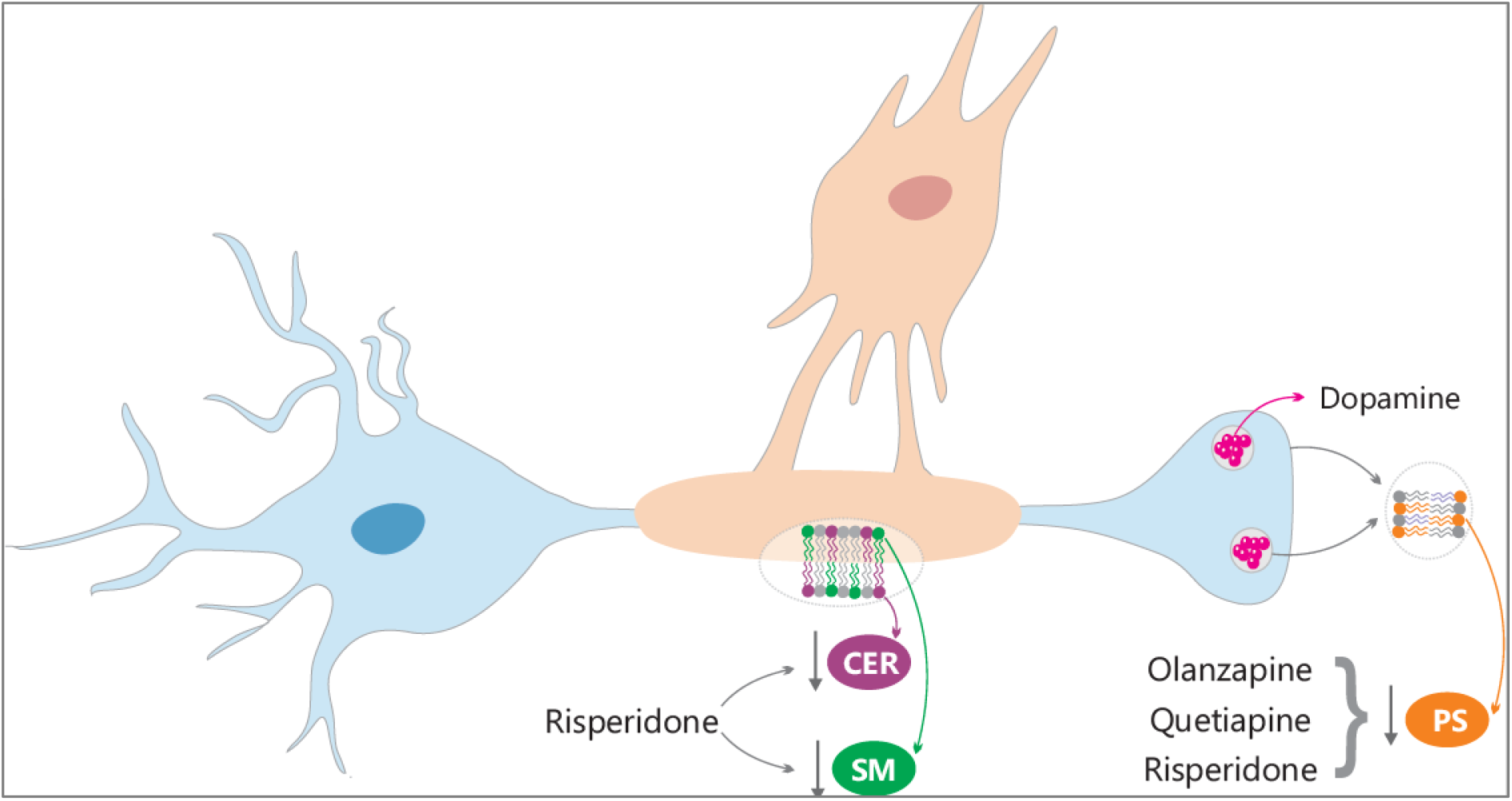
Possible mechanisms associated with poor response for antipsychotics. Antipsychotic-induced decrease in SM, CER, and PS lipids in the terminal neuron button and oligodendrocytes. Legend: CER, ceramides; PS, phosphatidylserine; SM, sphingomyelin.

Among all lipid classes, we noted that PCs were largely affected in our study. Accordingly, Leppik and colleagues (Leppik et al. 2019) found a decrease in 16 PCs in antipsychotic-näive first-episode psychosis. After 7 months, the authors reported that some PCs were up-regulated by antipsychotics, and these effects may be associated with modulation of the inflammatory pathways. Herein, we observed increased PC (38:4) in good responders treated with olanzapine and risperidone. However, this change was not specific; risperidone affected in opposite manner - decreased and increased - others lipids from PC class were observed in (PC40:8, and PC40:7) poor and (PC40:5, PC-O-42:1, PC38:5, PC-O-32:2, and PC-P-38:1) good responders by risperidone. Studies have shown that PC may be involved in schizophrenia pathobiology and antipsychotic mechanisms. Miller et al. (2012) found changes in PC in some brain regions of schizophrenia patients (Miller et al., 2012). Another study using ^31^P magnetic resonance spectroscopy found changes in PC in cerebrospinal fluid (Weber-Fahr et al., 2013), in the anterior cingulate, and left thalamus (Miller et al., 2012) of schizophrenia patients compared to healthy subjects. Nevertheless, opposite effects on PC levels were observed in this study, raising doubts about our conclusions of the role of these lipid class in antipsychotic treatment response.

The phosphatidylinositol (PI38:4) was up-regulated in good responders for the quetiapine treatment group and it was not affected by the others treatments. *Postmortem* studies found decreased levels of PI in the prefrontal cortex of schizophrenia patients compared to healthy subjects (Matsumoto et al., 2017). Additionally, this study found different pattern on the distribution of PI between gray matter and white matter in the prefrontal cortex of patients. These findings point out to possible disturbances on signaling in the prefrontal cortex, since PI in membrane cells can provide secondary messengers, such as inositol trisphosphate (IP3) and diacylglycerol (DAG), for several pathways. Thus, the increase in PI promoted by quetiapine may be associated with better clinical outcomes in schizophrenia, as observed in our study.

Herein, we observed that lipids from phosphatidylethanolamine (PE22:0) were increased by the risperidone treatment in good responders. A study found the same trend in lipids from the PE class in the white matter of frontal lobe samples of schizophrenia patients (Ghosh et al., 2017), supporting a role of lipid dysfunction in white matter deficits. In contrast, another study did not find changes in PE in dorsolateral prefrontal cortex samples from schizophrenia patients (Beasley et al., 2017). Others findings have been reported in the literature; one study showed decreased levels of PE in blood samples from first episode patients compared to healthy controls (McEvoy et al., 2013). The PE levels were found lower in first-episode psychotic (FEP) and chronic patients compared to healthy subjects, and these effects appeared not to be related to antipsychotic treatment (Kaddurah-Daouk et al., 2007). Together, these findings reinforce the potential role of PE in schizophrenia pathobiology. However, antipsychotic treatment may also modulate PE levels. One study found decreased PE levels in patients compared to controls, and this change was reversed by risperidone and olanzapine treatment (Kaddurah-Daouk et al., 2007). Herein we showed that only risperidone promoted changes on PE class in good responders.

Another common effect was the increase in glycerophosphoglycerol (PG) observed in good responders to olanzapine (PG-O-38:2 and PG-P-38:1) and quetiapine (PG34:1) treatments. However, PG lipids were found to be increased and decreased in poor responders of the olanzapine and risperidone groups, respectively. This suggests overall levels of PG lipids may not be a suitable marker of response. To date, no study has already reported the changes specific to this class induced by antipsychotics or in schizophrenia patients at baseline.

In the olanzapine group, we observed that glycerophosphatidic acids (PA) were down- and up-regulated in poor and good responders, respectively. The PA class plays a role in maintaining membrane curvature, intracellular pathways, and it can be a precursor for other lipids such as 2-arachidonoylglycerol (2-AG) synthesis, an endocannabinoid involved in schizophrenia pathobiology (Muguruza et al., 2013).

One specific effect in poor responders from risperidone was a decrease in sphingomyelin (SM (d17:1/24:1), SM(d18:2/23:0)) - an important lipid in the brain. Study showed changes in the expression of genes associated with sphingolipid metabolism in the prefrontal cortex (Narayan et al., 2009) and skin (Smesny et al., 2013) from schizophrenia patients. In addition, a lipidomic study reported a decrease of SM in red blood cells (RBCs) membranes of schizophrenia patients compared to controls (Tessier et al., 2016). A *postmortem* study also found decreased SM in brain samples of schizophrenia patients (Schmitt et al., 2004). In contrast, Keshavan et al. found increased SM in RBCs of patients (Keshavan et al., 1993). Specifically, Ponizovsky et al. demonstrated that increased SM in RBC was correlated with the negative symptoms (Ponizovsky et al., 2001). Although the samples used herein and in other studies were different, the findings taken together reinforce the potential role of SM in schizophrenia pathobiology and response to treatment. In agreement, one study found decreased SM levels in serum of antipsychotic-näive FEP patients; and the level of SM was decreased after 7 months of antipsychotic treatment (Leppik et al., 2019). These findings suggest that SM might be a potential biomarker for FEP and antipsychotic response, as we have hypothesized here.

SM is the most abundant sphingolipid in the central nervous system and plays a key role in the myelin sheath (Capodivento et al., 2017; Falkai et al., 2016; Martins-de-Souza et al., 2009). Studies have shown disturbances in myelin sheathing and oligodendrocytes in schizophrenia (Capodivento et al., 2017; Falkai et al., 2016; Martins-de-Souza et al., 2009). We observed a decrease in the ceramide 1-phosphates (CerP(d18:1/18:0)) level and lactosylceramides (LarCer) in poor responders of risperidone treatment. These findings lead us to suggest a potential mechanism of risperidone on SM and ceramide metabolism. Together with SM, ceramides comprise one of the most abundant lipids of the myelin sheath (Schmitt et al., 2015). Therefore, we suggest that the decreased levels of SM and ceramide lipids induced by risperidone (Figure 3) may be related to poor clinical outcomes. Furthermore, this effect of risperidone must be elucidated to avoid the negative effects of treatment of schizophrenia subtypes with more severe myelin disturbances (Figure 3). However, another lipid ceramide phosphoinositols (PI-Cer 38:0) was found decreased on good responder of the same group treatment. Additionally, a *postmortem* study found an increase of ceramides in white matter of schizophrenia patients, and that levels were not affected by antipsychotics (Schwarz et al., 2008). Recently, another *postmortem* study showed a decrease in ceramide and increase in lactosylceramides levels on frontal cortex of patients (Wood, 2019). These opposite findings reinforce that ceramides may not be a potential biomarker of response. However, we demonstrated that antipsychotics affected ceramide levels, at least in the blood, which could be considered for some conditions. Additionally, we hypothesized that risperidone’s effects on these lipid classes may be associated with 5-HT7 receptors (Knight et al., 2009). Since studies have shown association of sphingolipids with 5-HT7 receptor binding (Sjögren and Svenningsson, 2007). To note, quetiapine and olanzapine appear not to share this 5-HT7 receptor antagonist property.

## 5. Conclusions

Consistent findings had reported the increased risk of metabolic disturbances by atypical antipsychotics. Here we observed that risperidone treatment for 6 weeks affected more classes of lipids compared to olanzapine and quetiapine. Our study may help psychiatrists to evaluate the treatment according to their risk for metabolic disorders based on the blood levels of key lipids. Our study has also shown that some classes of lipids were specifically modified in the poor or good responder groups, which indicates that these may be potential biomarkers for antipsychotic treatment response. Interestingly, PS levels seem to be a potential biomarker for poor response, since we observed the same effect for all antipsychotics investigated in this study. Therefore, other antipsychotic drugs, such as typical antipsychotics or clozapine, could be indicated for such patients. Moreover, changes in lipids that were observed in good responders may highlight new targets for the development of new approaches to treat schizophrenia or to refine current drugs; however, further clinical trials are needed to reinforce these findings. Interestingly, quetiapine treatment induced only minor changes in lipids, and this group had more poor responders compared to the risperidone and olanzapine treatment groups. This reinforces a putative role of lipids in the therapeutic response to antipsychotics.

## 6. Limitations

These findings are limited by the small sample size and the need for validation using different sample cohorts. The samples used in this study are a subset of a larger cohort, but we selected those which came from drug-naïve or drug-free donors, to avoid any influence of pre-existing medications at baseline. Despite the reduced number of samples – a limitation for generalizing the conclusions addressed in this research – the models were adequately validated for this dataset, and therefore, providing the sufficient statistical robustness for the conclusions. Additionally, the modelling approach developed here can be employed with more sample cohorts for generalization. Thus, future studies in this area are warranted, especially considering the critical effects that drug-related MetS can have on the lives of patients suffering from schizophrenia.

## 7. Contributions

DMS conceived, organized, and supervised all steps of the study. VA wrote the paper together with DMS. JS collected and provided blood plasma samples and contributed with paper writing. AA prepared plasma samples and ran the mass spectrometry experiments helped by AG and MM. GA performed all statistical analyses and contributed with paper writing. PG helped in data analyses. VA and GA contributed equally to this work. All authors had access to and approved the final version of the manuscript.

## Supplementary Information

**Table S1:** Patients demographics

**Table S2:** List of compounds associated to the good or poor response

